# Grid Cells, Border Cells and Discrete Complex Analysis

**DOI:** 10.1101/2023.05.06.539720

**Authors:** Yuri Dabaghian

## Abstract

We propose a mechanism enabling the appearance of border cells—neurons firing at the boundaries of the navigated enclosures. The approach is based on the recent discovery of discrete complex analysis on a triangular lattice, which allows constructing discrete epitomes of complex-analytic functions and making use of their inherent ability to attain maximal values at the boundaries of generic lattice domains. As it turns out, certain elements of the discrete-complex framework readily appear in the oscillatory models of grid cells. We demonstrate that these models can extend further, producing cells that increase their activity towards the frontiers of the navigated environments. We also construct a network model of neurons with border-bound firing that conforms with the oscillatory models.

## I. INTRODUCTION AND MOTIVATION

Spiking activity of spatially tuned neurons is believed to enable spatial cognition [1–3]. For example, rodent’s *place cells* ^1^ that fire in specific locations produce a qualitative map of the explored environment [4–8]; *head direction* cells that fire each at its preferred orientation of the animals’ head contribute directional information [9–11]; the *grid cells* that fire near vertexes of a planar triangular lattice are believed to provide a metric scale [12, 13] and the *border cells* highlight the boundaries of the navigated enclosures [14–16] (Fig. 1A).

**FIG. 1.**
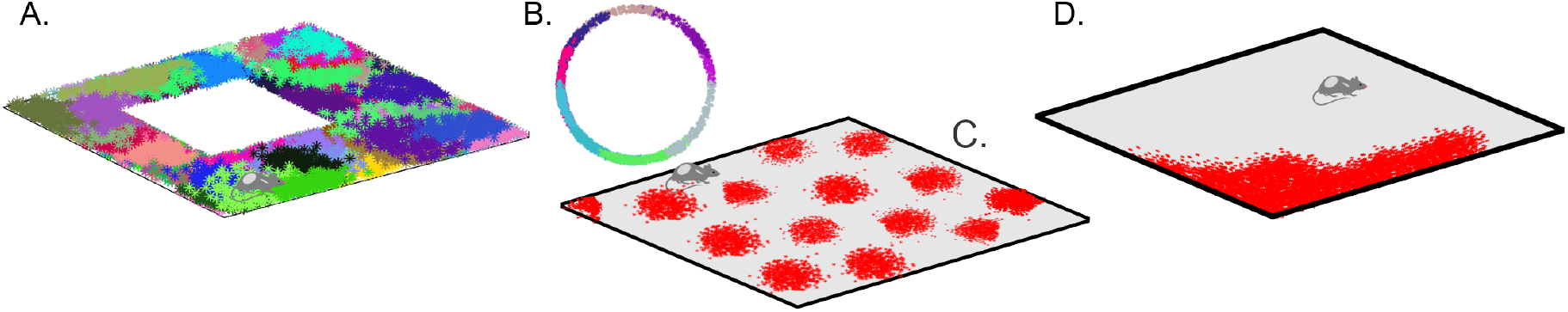
Spatial cells. (**A**). Spikes produced by place cells (dots of different colors) form distinct spatial clusters in the navigated environment, which highlight the preferred spiking domains—place fields. **B**. Head direction cells fire when the animals’ head is oriented at a particular angle with respect to cardinal directions, thus producing spike clusters in the circular space of planar directions. **C**. Spiking domains of the grid cells form a triangular lattice that tiles the ambient space. **D**. Boundary cells produce spikes along the border of the navigated enclosure.

A number of theoretical models aim to explain the machinery producing these spiking profiles, by exploiting suitable mathematical phenomena, e.g., attractor network dynamics [17–21], specific network architectures [21–24], the hexagonal symmetry of closely packed planar discs [25], constructive interference of symmetrically propagating waves [26–29] and so forth. In contrast, the ability of border cells to identify the frontiers of the explored environments was heretofore explained heuristically, as a certain “responsiveness” these neurons to the walls of the navigated arenas, achieved, conceivably, by integrating proprioceptive and sensory inputs [30–34]. However, since border cells are anatomically removed from sensory pathways, it is possible that their spiking may be produced through autonomous network mechanisms, rather than induced by external driving. From a computational perspective, such mechanisms may also hinge on a mathematical phenomenon that highlights the perimeters of spatial regions, a well known example of which is the *maximum principle*—the ability of certain functions, e.g., harmonic and complex-analytic functions, to attain maximal values at the boundaries of their domains [35].

The following study is motivated by a recent series of publications [36–39], which show that two-dimensional (2*D*) triangular lattices allow constructing a discrete counterpart of the Complex Analysis and defining real-valued, discrete epitomes of complex-analytic functions that obey the maximum principle. As it turns out, these structures allow modeling border cell activity, as discussed below.

The paper is organized as follows. Several key ideas of Discrete Complex Analysis (DCA) are outlined in Section II, following the exposition given in [36–39]. Section III discusses certain connections between elements of DCA and oscillatory interference models of grid cells [26–29], and offers a generalized framework for expanding these models to include border cell spiking patterns. In Section IV, elements of DCA are implemented in a schematic network model that produces border cell firing responses through endogenous activity, without using external parameters, such as animal’s speed or location. The results are briefly discussed in Section. V.

## II. APPROACH

### 1. Discrete complex analysis

Standard theory of complex variables is a calculus over complex numbers *z* = *x* + *iy* and their conjugates, 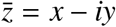, where *x* and *y* are the Cartesian coordinates in a Euclidean plane and *i* is the imaginary unit, *i*^2^ = −1 [35]. A generic complex function depends on both *z* and 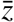; however, the main objects of the theory are the *analytic* (also called *holomorphic*) functions that depend only on *z, f* = *f* (*z*), and their *anti-analytic* (*anti-holomorphic*) counterparts, that depend only on 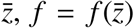. The defining property of these functions is that their derivatives over the “missing” variable vanish,

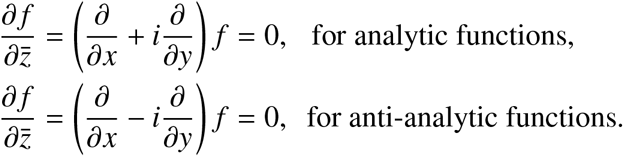

The Cauchy operator and its conjugate used above,

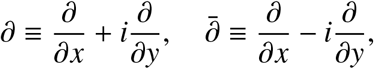

play key roles not only in complex analysis but also in geometry and applications. One of their properties is that they factorize the 2*D* Laplace operator, or the *Laplacian*,

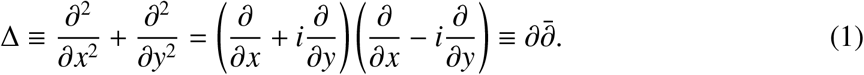

The factorization (1) is unique and necessarily involves complex numbers—think of the decomposition *x*^2^ +*y*^2^ = (*x*+*iy*)(*x*−*iy*) that is commonly used to motivate the transition from real to complex variables. Correspondingly, the phenomenon (1) takes place only on spaces that admit complex structure—orientable 2*D* surfaces. Furthermore, the factorization (1) can serve as a vantage point for defining the Cauchy operator and its conjugate: if a Laplacian admits the decomposition (1) in suitable coordinates, then the resulting curvilinear first-order operators 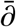 and *∂* will be the Cauchy operators of a complex-analytic structure on the corresponding manifolds.

A remarkable observation made in [36–39] is that the *discrete* Laplace operator on a 2*D* triangular lattice also is factorizable. Indeed, a generic discrete Laplacian on a graph or a lattice acts on the vertex-valued functions *f* (*v*) as

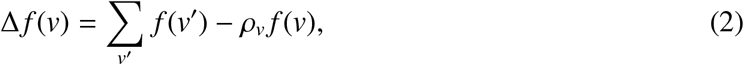

where the summation goes over all vertexes *v*^′^ linked to *v*, and ρ_*v*_ is the valency of *v* [40–42]. On a triangular lattice with vertexes marked by two integer indexes *m* and *n*, the Laplacian (2) becomes

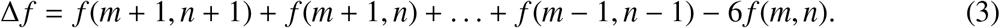

To obtain the required decomposition, let us define the operators τ_1_ and τ_2_ that shift the arguments of the vertexes functions,

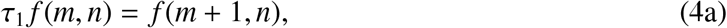

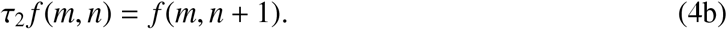

as shown on Fig. 2A. In terms of τ_1_ and τ_2_, the sum (3) becomes

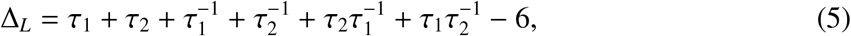

and factorizes into the product of two first-order operators

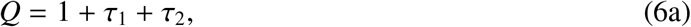

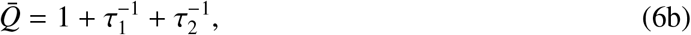

with an extra constant term,

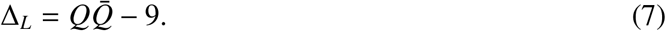

As shown in [36–39], this decomposition induces a DCA, in which the operator 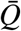 plays the role of the complex-conjugate derivative 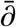. One can thus define the discrete-analytic lattice functions, *f* (*m, n*), as the ones that satisfy the relationship

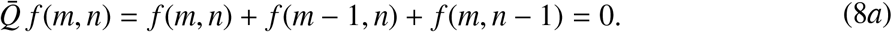

The *Q*-operator then acts as the discrete-analytic derivative,

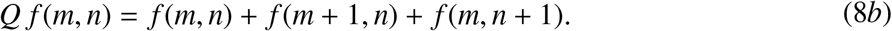

Geometrically, equations (8) can be illustrated by partitioning the lattice *V* with “black” and “white” triangles, in which each white triangle, △, shares sides with three black triangles, ▾, and vice versa (Fig.2B). According to (8*a*), the discrete analytic functions vanish over all the black triangles, which may be viewed as the lattice analogue of “*z*-but-not-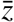 dependence of the conventional complex-analytic functions.

**FIG. 2.**
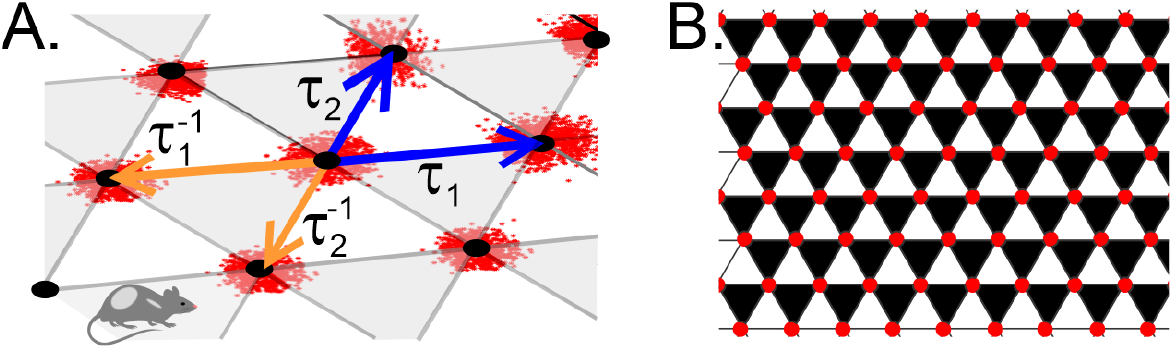
Lattice. **A**. The operators τ_1_ and τ_2_ (blue arrows), shift the argument of the lattice function forward along the basis directions, from (*m, n*) to (*m* + 1, *n*) and (*m, n* + 1) respectively. The inverse operators, 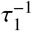 and 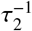 (orange arrows), shift the argument backwards, correspondingly to (*m* − 1, *n*) and (*m, n* − 1). **B**. The backwards shifts 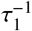 and 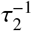 support the “black” triangles and forward shifts τ_1_ and τ_2_ span the complimentary set of “white” triangles. If a function satisfies the discrete-analyticity condition (8*a*), then its values over the black triangles vanish.

### 2. Properties of the discrete-analytic functions

largely parallel the familiar properties of their continuous counterparts, including the maximal principle that is used below to model the border cell spiking activity. However, there are also a few differences, the most striking of which is that the discrete-analytic functions are *real-valued*: indeed, the equation (8*a*) does not involve imaginary numbers and possesses real-valued solutions [36–39]. Thus, the discrete complex analysis is a real-valued combinatorial framework that may be implemented through neuronal computations^2^.

Another peculiarity is that DCA redefines the notion of a constant. Indeed, the constants *c* of the standard calculi are nullified by the derivatives, 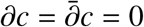. However, a quantity that assumes constant values on all vertexes, *f* (*m, n*) = *c*, is not nullified, but tripled by discrete derivative operators, 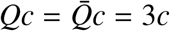. Hence, discrete-analytic constants *h* must be derived from the equations

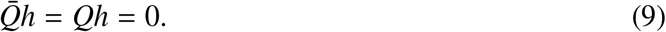

Somewhat surprisingly, the basic solutions of (9) have the form

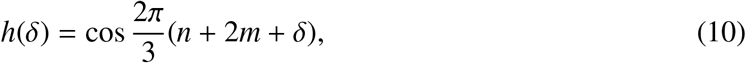

where *δ* is a phase parameter (Fig. 3A). Formula (10) can be viewed as a discrete analogue of the complex phase *e*^*iδ*^; the “prime” constants 1 and *i* then correspond to

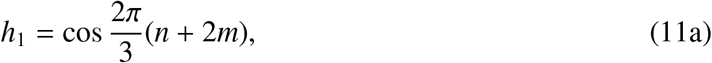

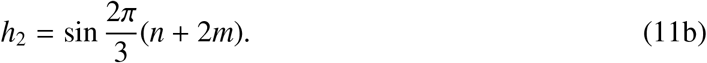

Note that, in contrast with their familiar counterparts, the “constants” (10) and (11) alternate from vertex to vertex, assuming a few discrete values, *h*_1_ = {−0.5, 1} and 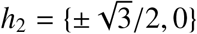.

**FIG. 3.**
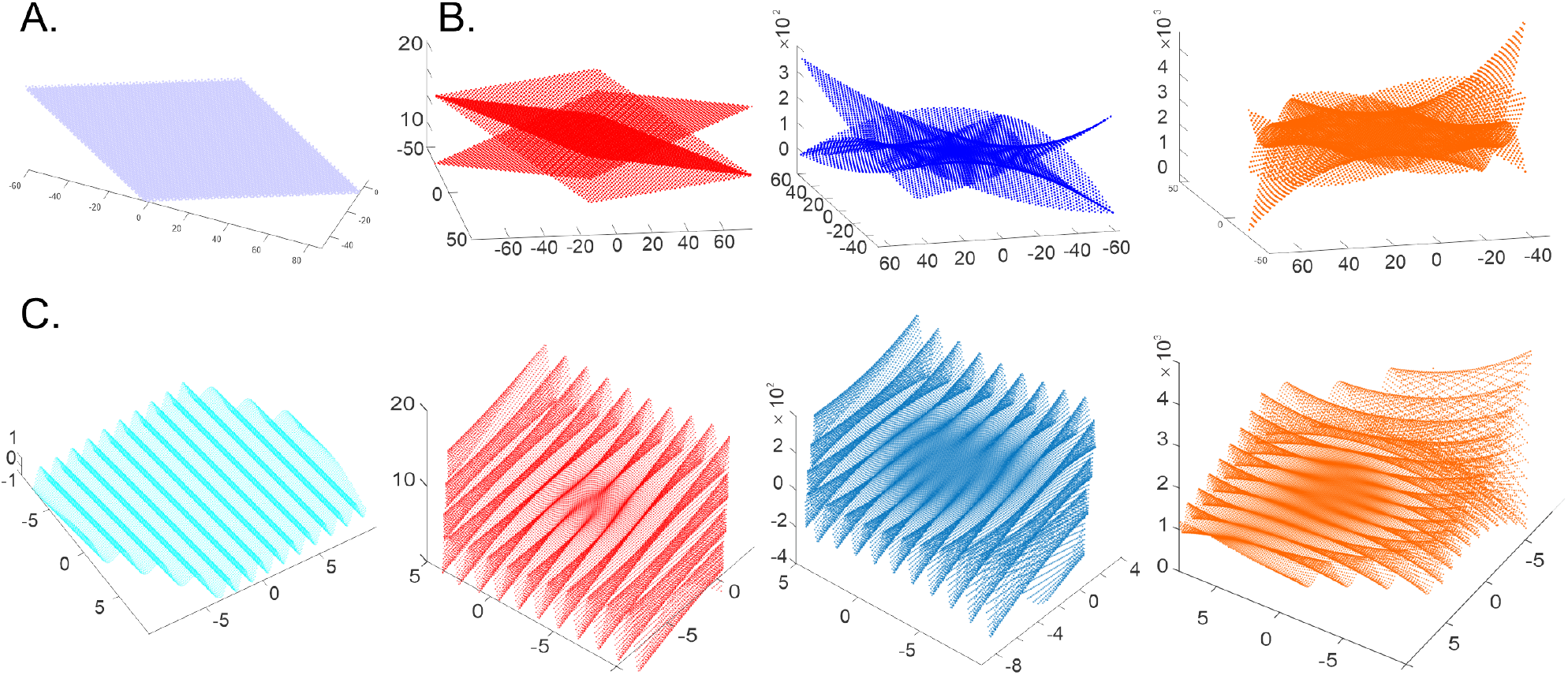
Discrete-analytic polynomials. **A**. 0^th^ order polynomials are the holomorphic constants that assume a small set of discrete values *h*_1_ = {−0.5, 1} and 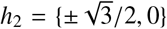. **B**. The discrete-holomorphic polynomials *P*_1_(*m, n*), *P*_2_(*m, n*) and *P*_3_(*m, n*) grow outwards (linearly, parabolically and cubically) as the lattice indexes increase. **C**. The spatially-refined discrete polynomials produce undulatory shapes scaffolded by their discrete counterparts: shown are the undulating holomorphic wave *h*_1_(*x, y*) and the polynomials *P*_1_(*x*_1_, *x*_2_), *P*_2_(*x*_1_, *x*_2_) and *P*_3_(*x*_1_, *x*_2_) that grow towards the boundary of the enclosed Euclidean domain.

The third distinct property concerns Taylor-expansions: in contrast with the continuous case, a generic discrete-holomorphic function *f* (*m, n*) over a finite l attice d omain can be represented exactly by finite series, i.e., one can write

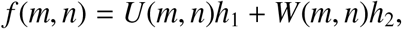

where *U*(*m, n*) and *W*(*m, n*) are polynomials. The order of such polynomials generally grows with the size of the lattice domain, which allows keeping the above expansion exact.

Explicit examples of the first, second and third-order discrete-analytic polynomials are

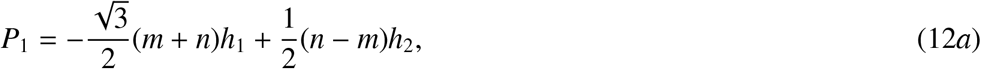

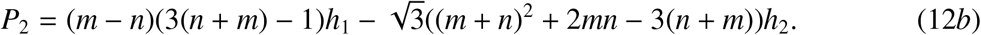

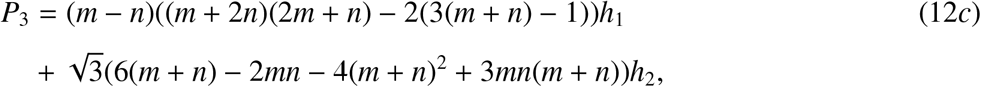

illustrated on Fig. 3B, C and D. It can be verified by direct substitution ^3^ that the operator 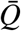 nullifies each polynomial, whereas *Q* lowers their order, *QP*_1_ = *h*_1_, *QP*_2_ ∝ *P*_1_ and *QP*_3_ ∝ *P*_2_, just as 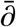 would nullify polynomials of *z*, and *∂* would lower their order, *∂p*_*r*_(*z*) ∝ *p*_*r*−1_(*z*). In general, there are 2(*r* + 1) basic discrete-analytic polynomials of order *r*, which corresponds to 2(*r* + 1) basic complex *r*^th^-order complex polynomials [36–39].

### 3. Spatial fine-graining

Discrete functions defined over the lattice vertexes give rise to finer-grained spatial structures. Given two basis vectors

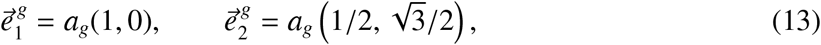

in the Euclidean plane, consider a lattice generated by integer translations,

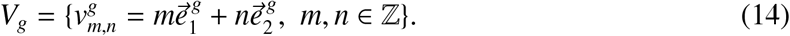

Such embedding allows extending the discrete argument of a vertex function, *f* (*m, n*), to a function of Euclidean coordinates, *f* (*x*_1_, *x*_2_), by replacing the integer arguments (*m, n*) with pairs of reals (*x*_1_, *x*_2_). For example, the discrete-holomorphic constant (11a) yields a continuous “holomorphic wave” with wavelength ∝ *a*_*g*_,

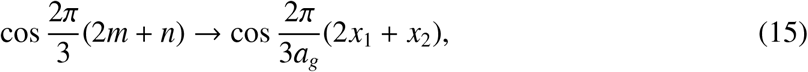

propagating in the direction 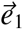 (Fig. 3C). Conversely, using

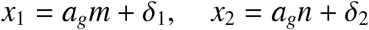

in the real-valued functions with sufficiently low spatial frequency (less than 2π/*a*_*g*_) restores the dependence upon the lattice indexes and produces a continuous phase *δ* that contains fractional remainders,

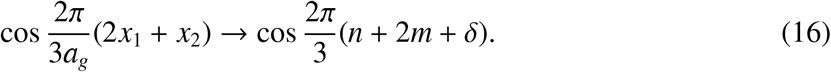

The latter form allows acting with the operators *Q* and 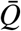 on the regular coordinate functions and placing the results into the context of DCA.

## III. OSCILLATORY GRID CELL MODELS

Surprisingly, discrete-analytic structures are manifested in the existing models of grid cell activity, e.g., in the oscillatory interference models that derive the observed grid field patterns from the dynamics of the membrane potential,

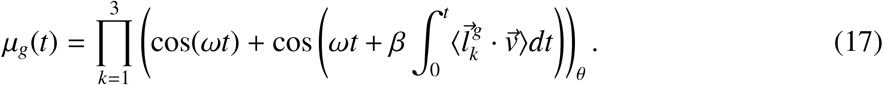

Here *t* is time, β is a scale parameter, 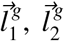 and 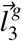 are the three symmetric wave vectors, 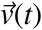 is the velocity, and ω ≈ 8 Hz is the frequency of synchronized extracellular field’s oscillations. The index “θ” refers to the firing threshold [26–29]. Due to the symmetry, the waves interfere constructively at the vertexes of a triangular lattice with basis vectors 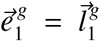 and 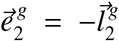, centered at the firing fields ^4^ (Fig. 1C, 4A).

**FIG. 4.**
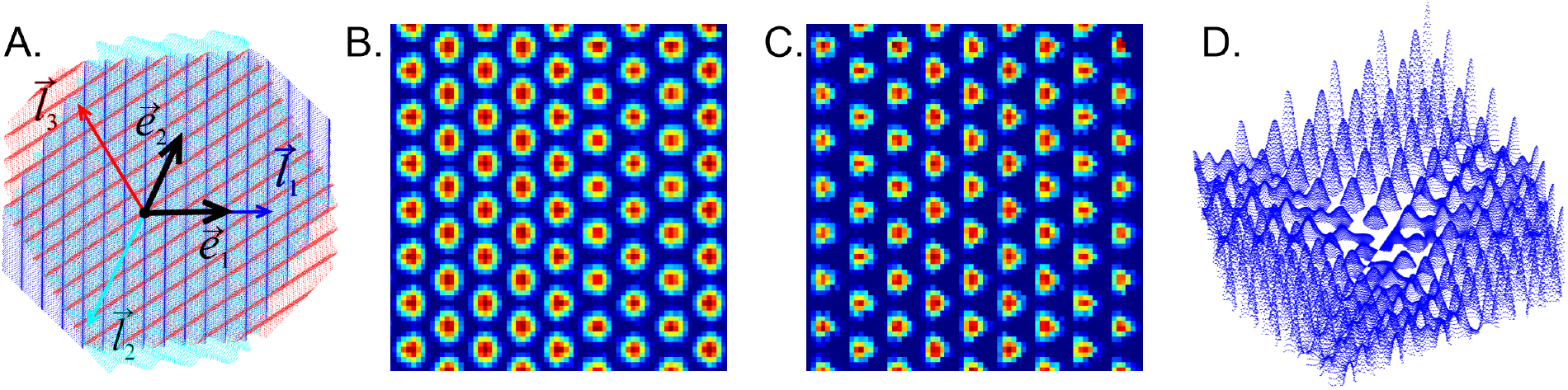
Oscillatory interference model. **A**. Superposition of three discrete-holomorphic waves propagating in three symmetric directions specified by the three wave vectors 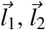 and 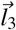. The basis lattice directions 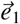 and 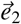 are shown in black. Constructive interference occurs at the vertexes of a triangular lattice, highlighted by the amplitude (18). **B**. The grid cell firing amplitude, *A*_*g*_, formula (19), reproduces the familiar grid cell layout. **C**. The complementary “conjugate” amplitude, 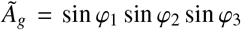. **D**. The second order discrete-holomorphic *grid-polynomial* (21*b*), defined over the grid fields, compare with the third panel on Figs. 3C.

To link *µ*_*g*_(*t*) to DCA, let us rewrite the time integrals in (17) as integrals along the trajectory,

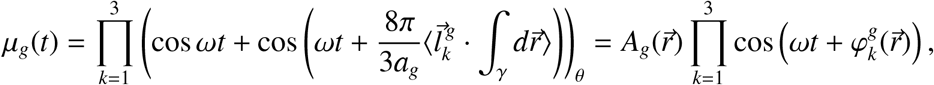

where 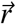 is the position vector, 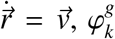 are the oscillatory phases and 8π/3*a*_*g*_ = *β*. The timeindependent factor defines the spatial amplitude of the membrane potential,

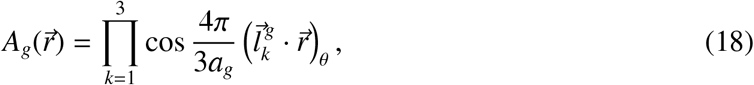

and produces the familiar spatial pattern of grid fields, brought about by the constructive interference of the contributing waves (Fig. 4B). Next, given the rat’s position in the lattice basis, 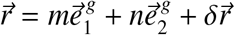 and using 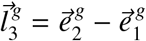, yields

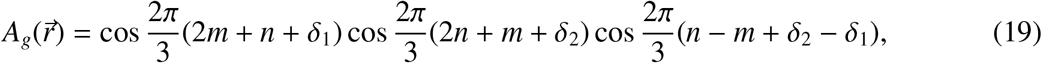

where 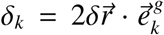 are the remainder phases. Curiously, each multiplier in (19) is a discrete-holomorphic constant: the second coincides with (10), the first can be obtained from (10) by reindexing, *m* ? *n*, and the last is produced by an index shift, *n* → *n* − 3*m*. Even more surprisingly, the full product (19), adjusted by a constant reference value 1/4, is also nullified by the discrete Cauchy operators,

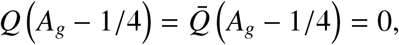

which means that the amplitude of grid cells’ firing (18) is, in fact, a basic DCA object—a finegrained discrete-holomorphic constant that functionally highlights the lattice of firing fields. The neurons that respond to grid cell outputs can hence be viewed as functions on that lattice, which includes discrete-holomorphic functions used for modeling border cells. Furthermore, the necessary elements of the DCA can be constructed independently within the oscillatory model, as discussed below.

## IV. BORDER CELLS

### Oscillatory model

of the grid cells can be generalized to simulate border cells’ activity by replacing the constant membrane potential (17) with suitable discrete-holomorphic functions obeying the maximum principle. The resulting firing rate will then grow towards the boundary of the navigated environment ℰ and produce the characteristic border cell firing patterns.

A simple implementation of this idea can be achieved using the discrete-analytic polynomials (12), by replacing the combinations

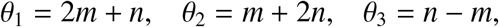

with the phases appearing in (18),

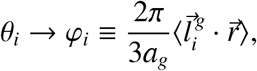

that represent dendritic inputs into the postsynaptic cell [43]. The resulting fine-grained discreteholomorphic polynomials are then

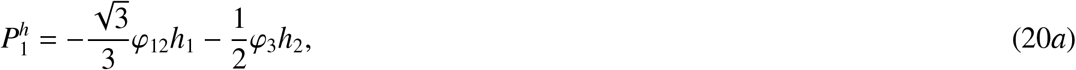

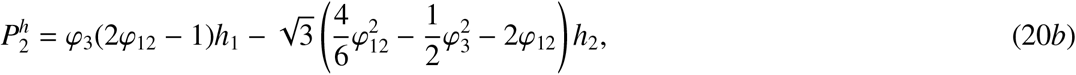

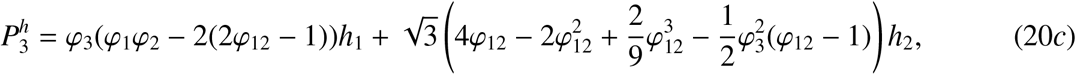

where φ_12_ is a short notation for (φ_1_ + φ_2_)/2 and the waves *h*_1_, *h*_2_ in (12) can be steered along any of the symmetric directions, 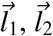 or 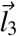.

Physiologically, it is possible^5^ that border cell activity is gated by inputs from the grid cells [44–49]. This mechanism can be modeled by replacing the “undulating” holomorphic constants *h*_1_ and *h*_2_ in (12) with the grid cell firing amplitudes, *A*_*g*_ and the complementary combination of holomorphic sine waves 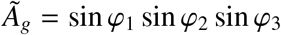 (Fig. 4C), which yields *grid polynomials*, e.g.,

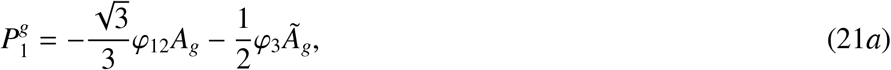

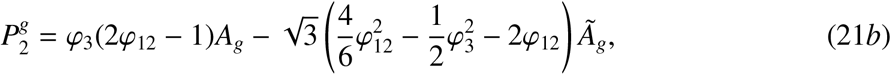

defined explicitly over the grid field lattice (Fig. 4D). By direct verification, both sets of polynomials (20) and (21) are discrete-analytic functions that obey the maximum principle and can hence serve as building blocks for producing generic membrane potentials accumulating towards the boundaries of the navigated enclosures.

As mentioned above, the individual φ-terms in (20) and (21) may be physiologically interpreted as the inputs received through linear or nonlinear synapses. Since the second- and the third-order nonlinear synapses are discussed in the literature [50–60], we used combinations of 5-10 polynomials of the orders *r*_*i*_ = 1, 2, 3,

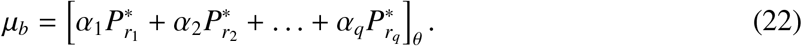

Here the 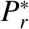 represent either harmonic (20) or the grid polynomials (21), the coefficients α_*i*_ define the magnitude of each addend, and the θ subscript indicates the threshold. In the simulations, the values α_*i*_ were selected randomly, while the threshold grew according to the size of the environment and the order of the contributing polynomials, 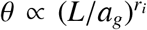. The resulting firing maps are illustrated on Fig. 5A,B. Expectedly, since all contributing polynomials in (22) grow towards the boundaries of the available lattice domain, all simulated border cells fire along the frontiers of the navigated enclosure.

**FIG. 5.**
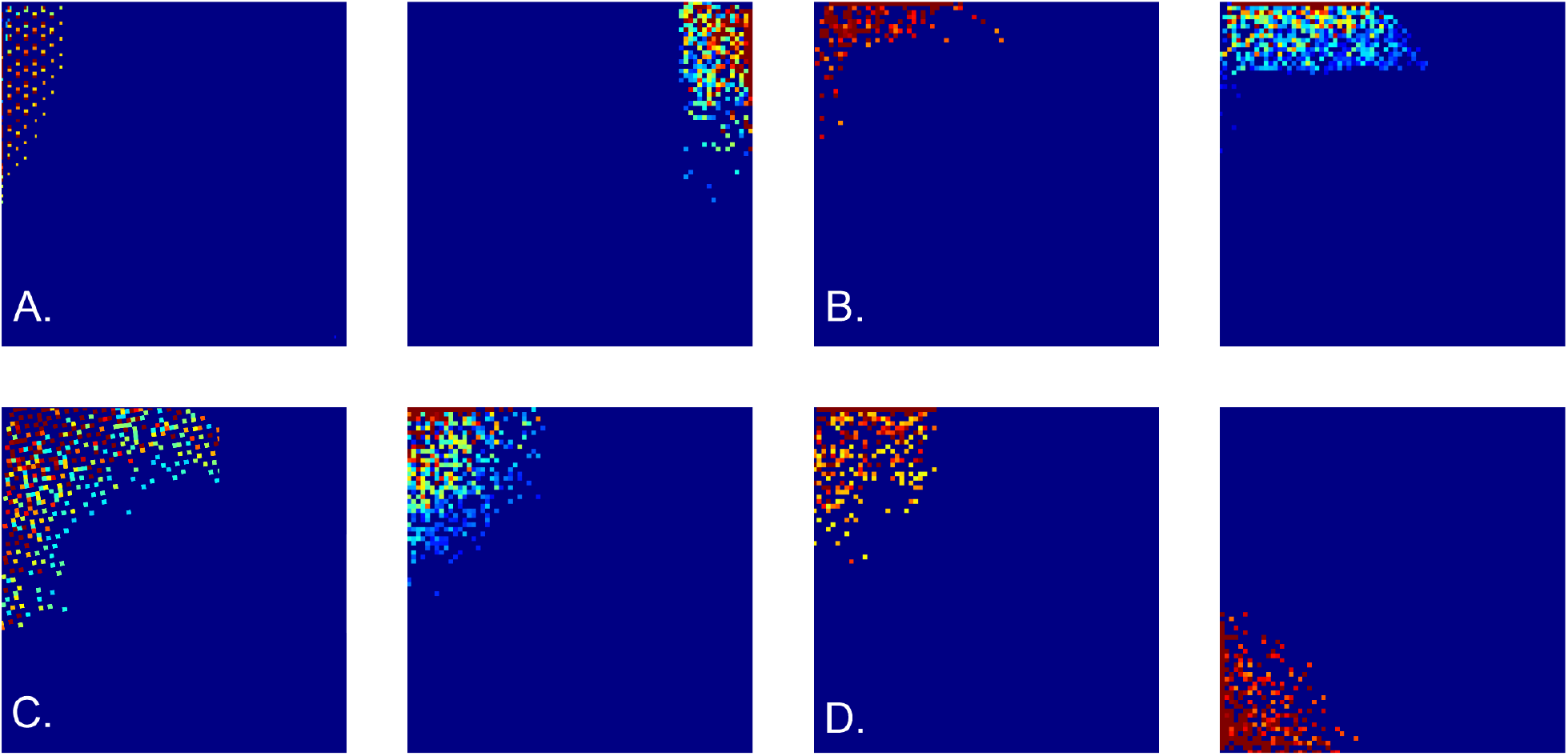
Border cell firing patterns. Examples of simulated border cell firing fields in a square 6 × 6 m environment, obtained as combinations of first, second, and third order grid polynomials. **A**. Firing fields obtained using “undulating” discrete-holomorphic polynomials (12). **B**. Examples of the firing fields obtained using combinations of grid-holomorphic polynomials (21). **C**. Firing fields of noise-perturbed membrane potentials, for *ε* = 0.2 (left panel) and *ε* = 0.3 (right panel). **D**. Firing fields obtained using the schematic network model.

Importantly, these outcomes are robust with respect to stochastic variations: disturbing the phases φ_*i*_ of the holomorphic polynomials with a noise term, *εξ*, where *ξ* is a random variable uniformly distributed over [0, 2π] and *ε* controls its amplitude, does not qualitatively alter the resulting spatial patterns for *ε* ≤ 0.5 or more (Fig. 5C).

### Schematic network model

Defining the membrane potentials as functions of speed and coordinates used, e.g., in (17) helps linking the geometry of the observed environment to the underlying neuronal computations. However, modeling the brain’s own representation of the ambient environment requires using intrinsic representation of spatial information, a key role in which is played by hippocampal place cells, *c*_*i*_, and the postsubicular^6^ head direction cells, *h*_*i*_ [3, 9]. The computational units enabling this representation are the functionally interconnected cell groups

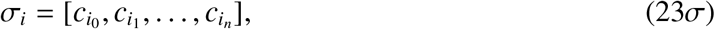

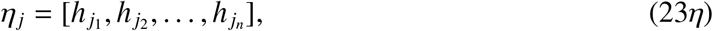

which highlight, respectively, basic locations 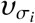 and angular domains 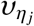 [61–65]. A number of studies have demonstrated that the assemblies (23) encode the animal’s ongoing position, the shape of trajectory and even its planned and recalled navigational routes [66–72]. By the same principle, place cell assemblies that fire over the grid fields 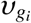, can provide their hippocampal representation: a combination 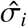 of σ-assemblies whose constituent cells exhibit coactivity with a grid cell *g* and each other defines a vertex of grid cell activity,

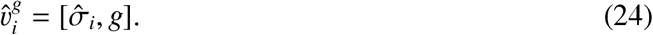

In the following, the superscript “*g*” will be suppressed in describing single grid cell activity and used only to distinguish contributions from different grid cells.

The hexagonal order on the vertexes (24) is established by concomitant activity of select groups of head direction assemblies, 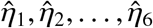, that activate on the runs between pairs of neighboring grid fields, e.g., *υ*_*i*_ and *υ* _*j*_, thus defining the *spiking edges* between 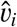 and 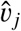,

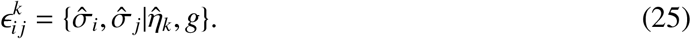

Together, the vertexes (24) and the edges (25) can be viewed as elements of a *spike-lattice* 𝒱_*g*_, by which the grid field lattice is embedded in the cognitive map [73]. Using 𝒱_*g*_ allows constructing a self-contained phenomenological network model of border cells that does not involve “tagging” the neuronal activity by externally observed characteristics, such as the rat’s speed or Euclidean coordinates.

Suppose that a cell *b* with membrane potential *µ*_*b*_ receives input from a group of persistently firing head direction assemblies 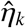, over a period when grid cell *g* becomes active, then shuts down, and then restarts its activity again^7^. If these consecutive activations are induced over adjacent vertexes 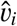 and 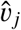, then the corresponding change of the membrane potential can be interpreted as the change of the spike-lattice function 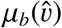 along the edge *ϵ*_*i j*_ between them,

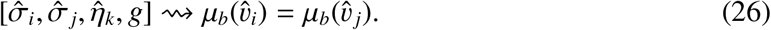

On the other hand, the transformation (26) can be described as the action of a *spike-lattice shift operator* 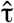 on *µ*_*b*_,

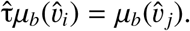

In particular, changes induced by the head direction assemblies 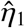 and 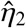 (ordered as on Fig. 2A) can be identified with the shift operators acting “forward” along the basic lattice directions,

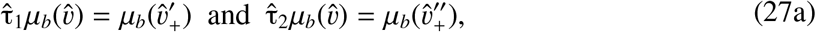

while the “opposite” assemblies 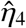 and 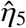 induce backward transformation,

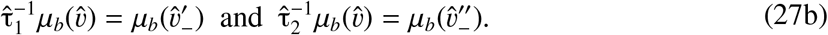

The appearance of spiking analogues of the shift operators τ_1_ and τ_2_ associated with grid cells opens a possibility of implementing the key DCA structures neuronally. However, a principal challenge in this approach is that the series of inputs received along a particular trajectory may not concur with the lattice structure of the underlying grid fields. Indeed, consider the membrane potential at the initial spiking vertex 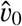,

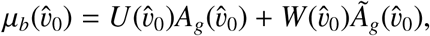

from where the animal continues to move along a trajectory γ, producing a series of postsynaptic changes described by a sequence of 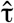-shifts,

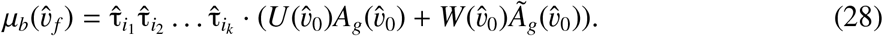

If the net membrane potential (28) does not depend on the order in which the individual inputs arrive, the 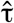-operators commute^8^. Thus, the value accrued at the final vertex 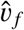 is

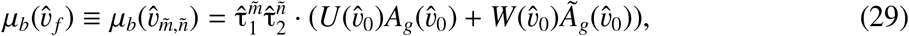

where the integers 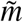 and 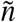 mark how many times 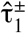 and 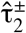 were triggered along the way.

Note however, that a generic trajectory γ may not pass through the fields of a given cell *g* in complete sequence: some fields are visited, others are occasionally missed (Fig. 6A). As a result, the “empirical” 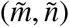-indexing appearing in (29) may not conform with the original (*m, n*)- indexing of the full grid field set, which moots the possibility of interpreting the argument of *µ*_*b*_ in terms of the underlying lattice (14). However, it can be shown that, within physiological parameter range, there typically exists a special class of “percolating” paths—those that run through the firing fields of a given grid cell in contiguous sequence, without omissions (see [73] and Fig. 6A). Such paths induce series of conjoint spiking edges,

**FIG. 6.**
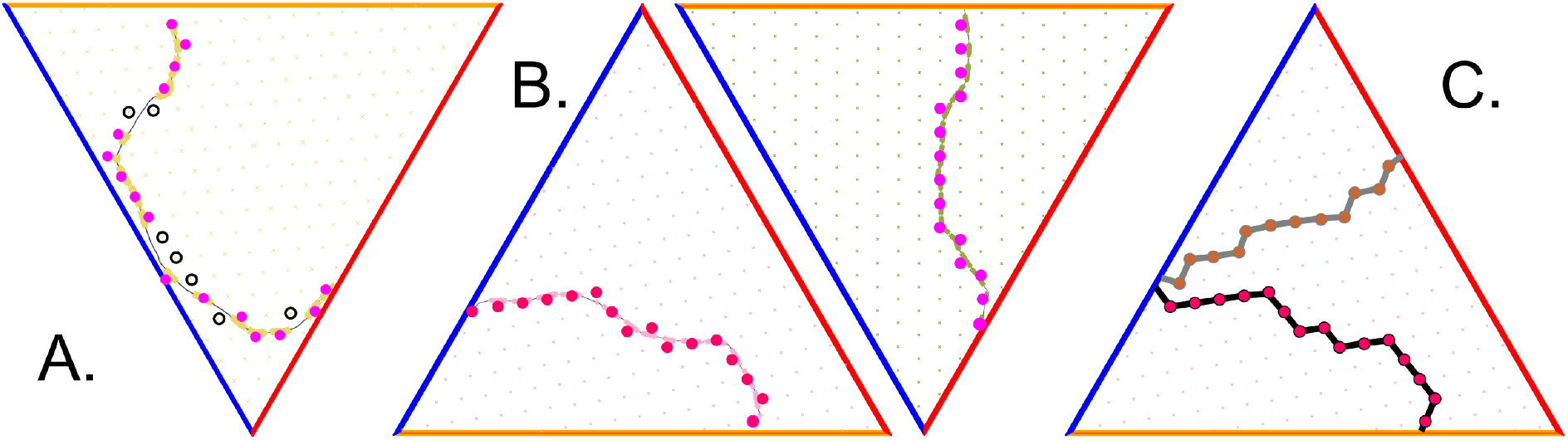
Grid cell percolation and border cell firing patterns. **A**. Generic path is non-percolative: the vertexes that correspond to the “percolated” firing fields—the ones over which the grid cell has produced at least one spike—are marked by pink, while vertexes corresponding to the fields that did not respond are marked by black. **B**. Two examples of percolating trajectories, along which the spiking occurred at each vertex, without omissions. **C**. Two examples of lattice paths induced by two percolating trajectories.

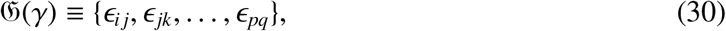

that serve as lattice representations of the animals’ moves (Fig. 6C, [73]). The increments of the postsynaptic membrane potential (29) acquired along the link series (30) are, by design, compatible with the lattice indexing and hence allow constructing consistent lattice functions over an extended lattice domains [73]. The subsequent development of the model will therefore be based on percolating paths only.

Constructing a membrane potential (29) by applying spiking 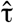-operators along the percolated paths requires knowing how these operators act on discrete-holomorphic constants and polynomials, which can be established as follows. First, the response of the spike-lattice counterparts of holomorphic constants *h*_1_, *h*_2_, and of their grid analogues, 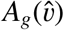 and 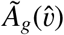, to 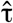-shifts (27), can be implemented according to how the corresponding original, index-dependent expressions (11) and (19) respond to the τ-operators, e.g.,

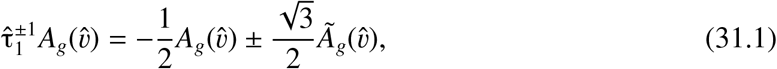

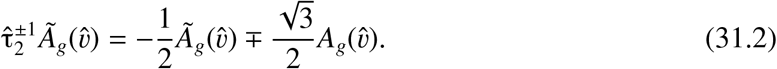

One can then use the expressions (31) along with (27) as the rules defining how the 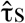 act on the spike-lattice 𝒱_*g*_, and thus deduce how the “spiking” Cauchy operator 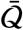 acts on generic membrane potentials,

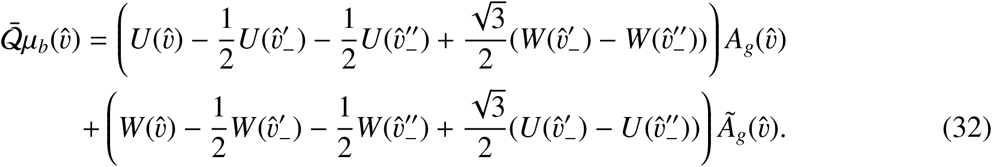

To satisfy the discrete analyticity condition, 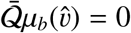, the coefficients in front of the holomorphic constants in (32) must vanish at each spike-vertex 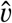. The simplest solution to this requirement is provided by the functions that acquire constant increments over the vertex shifts,

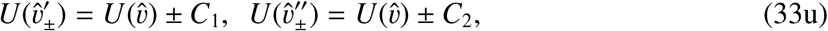

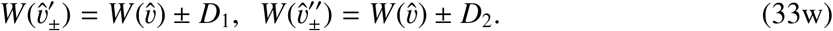

By direct verification, the equation 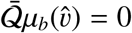 is satisfied identically if

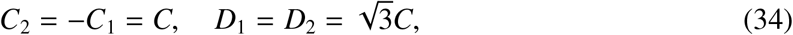

where *C* represents vertex-independent additive synaptic input. Thus, if the specific synaptic responses to each of the τs are defined by (34), then the net accumulated postsynaptic membrane potential is

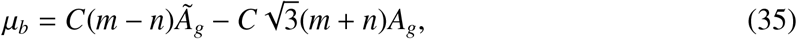

which matches the linear discrete-holomorphic polynomial (21*a*) and clarifies how such potential may emerge through synaptic integration. For the nonlinear membrane potentials described by higher-order polynomials, the shifting rules can be obtained by analogy with (33), by requiring that the shifted values are described by lower-order polynomials, e.g., by linear increments to the shifted second-order polynomials,

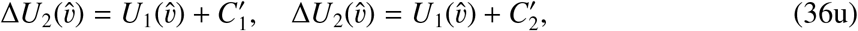

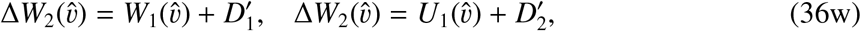

and so forth. The results then produce second and third order expressions of the type (20*b*) and (21*b*), which combine according to (22) and yield build border cell firing patterns as illustrated on Fig 5D.

## V. DISCUSSION

A number of computational models aim to explain the origins of the triangular spatial pattern of the grid cells’ spiking activity and the contribution that these cells make into enabling spatial cognition [20]. It is believed that the regular grid firing patterns allow establishing global metric scale in the navigated environment [13] and may produce a spatial location code [77–79]. The model discussed above shows that the neuronal mechanisms producing hexagonal layout of the firing fields may enable yet another mathematical phenomenon—a discrete complex structure. Although the whole structure is implemented via real-valued computations, it captures all the key attributes of the conventional theory of complex variables. In particular, the discrete-analytic functions defined in DCA framework obey the maximum principle—a property that may be used to model neurons with firing responses tuned to the boundaries of the navigated environments.

Surprisingly, basic elements of DCA are manifested in several existing models of grid cells. As discussed above, the interfering waves of the oscillatory models, which may be viewed either as representation of physiological rhythms, or as formal spatiotemporal components of the membrane potential’s decomposition, can be described as spatially fine-grained discrete holomorphic constants. Their net interference pattern, that defines the grid cells’ firing amplitude, also amounts to a discrete-holomorphic “grid” constant, highlighting a triangular lattice. Incorporating higher-order polynomials (and hence generic discrete-holomorphic functions) into the model allows simulating border cell spiking, which emphasizes affinity of the two firing mechanisms. The latter may explain why these cells are anatomically intermingled—in electrophysiological recordings, both cell types are often detected on the same tetrode. According to the model, physiologically similar neurons may be wired to perform synaptic integrations of different orders, and may, conceivably, swap their firing types in response to synaptic or structural plasticity changes.

The DCA approach can be also be used to produce self-contained network models that do not require phenomenological inputs, i.e., do not reference speed, coordinates, grid field positions, *ad hoc* lattice indexes (*m, n*) or other externally observed tags of neuronal activity. On the contrary, it becomes possible to render certain abstract DCA structures via autonomous network computations. For example, the Cauchy operators and the lattice (14) underlying the grid field layouts are induced using the “spiking” analogues of the τ-operators (4),

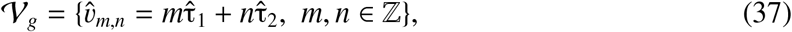

with vertex indexes derived from counting synaptic inputs of the grid, head direction and place cells along the percolated paths. In this context, the standard procedure of constructing grid fields 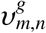 (Fig. 1C), by attributing (*x*_1_, *x*_2_) coordinates to spikes according to the rat’s ongoing location, can be viewed as a mapping from the vertexes of the spike lattice (37) into regions in the navigated environment,

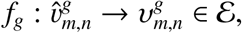

centered at the vertexes of the grid field lattice *V*_*g*_ [10, 80]. Zero holonomy property of the discrete Cauchy operators discussed in [36, 37] (see also [81]) ensures that the (*m, n*) values attained at a particular vertex do not depend on the percolating paths leading to a vertex, but only on the vertex itself, which ensures consistency of the construction. The discrete-complex structure can thus be viewed as an intrinsic network property, that may be implemented using different synaptic architectures, e.g., the continuous attractor models. An implication of this property is that the grid cells should be expected to produce planar, rather than voluminous firing fields, in order to implement the Cauchy decomposition (7) attainable only on 2*D* hexagonal lattices—a prediction that agrees with both experimental [82–86] and theoretical [87–90] studies.

As a concluding comment, the DCA framework currently does not offer a direct geometric interpretation of the discrete-holomorphic mappings [36–38]. An independently developed notion of discrete conformal transformations, based on rearrangements of regular circle packings in planar domains [91–94] may therefore offer a complementary venue for establishing correspondences between network activity and discrete-complexity. Several recent experimental [95–101] and theoretical [102–106] studies suggest that conformal transformations of the navigated spaces may induce compensatory discrete-conformal transformations of the grid field maps, similar to how the hippocampal place cells tend to preserve coactivity patterns in morphing environments [5, 7, 8]. If the latter is verified experimentally, it can be argued that the grid cell inputs constrain the hippocampal topological map [7, 8], to a net conformal map of the navigated space.

## Acknowledgments

Work was supported by NSF grant 1901338 and NIH grant R01NS110806.

## Footnotes

1 Throughout the text, terminological definitions and semantic highlights are given in *italics*.

2 DCA can also be constructed over the complex numbers: the corresponding theory then yields the standard Complex Analysis in the limit when the lattice side vanishes [36–39].

3 The operators *Q* and 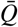 generally do not distribute according to the Leibniz rile, e.g., 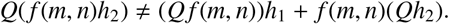

4 The model [25] uses sum of three waves for capturing analogous interference effect.

5 Currently, the synaptic organization of the border cell network is debated.

6 Head direction cells are also found in few other brain regions [9].

7 For a physiological discussion, see [73–76].

8 As do their τ-counterparts [36–39].

